# Single nucleotide polymorphisms of the *c-MYC* gene’s relationship with formation of Burkitt’s lymphoma using bioinformatics analysis

**DOI:** 10.1101/450783

**Authors:** Anfal Osama Mohamed Sati, Weaam Anwer Osman, Enas Abdalla Mohammed Ahmedon, Safa Hamed Elneel Yousif, Enas Dawoud Khairi, Alaa Ibrahim Mohammed Hassan, Moshtaha Ali Ibrahim Elsammani, Mohamed Ahmed Salih

## Abstract

Burkitt’s lymphoma (BL) is an aggressive form of non-Hodgkin lymphoma, originates from germinal center B cells, MYC gene (MIM ID 190080) is an important proto-oncogene transcriptional factor encoding a nuclear phosphoprotein for central cellular processes. Dysregulated expression or function of c-MYC is one of the most common abnormalities in BL. This study focused on the investigation of the possible role of single nucleotide polymorphisms (SNPs) in MYC gene associated with formation of BL.

MYC SNPs were obtained from NCBI database. SNPs in the coding region that are non-synonymous (nsSNPs) were analysed by multiple programs such as SIFT, Polyphen2, SNPs&GO, PHD-SNP and I-mutant. In this study, a total of 286 Homo sapiens SNPs were found. Roughly, forty-eight of them were deleterious and were furtherly investigated.

Eight SNPs were considered most disease causing [rs4645959 (N26S), rs4645959 (N25S), rs141095253 (P396L), rs141095253 (P397L), rs150308400 (C233Y), rs150308400 (C147Y), rs150308400 (C147Y), rs150308400 (C148Y)] according to the four softwares used. Two of which have not been reported previously [rs4645959 (N25S), rs141095253 (P396L)]. SNPs analysis helps is a diagnostic marker which helps in diagnosing and consequently, finding therapeutics for clinical diseases. This is through SNPs genotyping arrays and other techniques. Thus, it is highly recommended to confirm the findings in this study in vivo and in vitro.

## Introduction

Burkitt’s lymphoma (BL) is an aggressive form of non-Hodgkin lymphoma, originates from germinal center B cells ^(1-6)^. It accounts for 30 to 50% of lymphomas in children and for 1 to 2% in adults^(2,5-8)^. The WHO classifies Burkitt’s lymphoma into three clinical variants: endemic, sporadic (the Predominant type found in non-malarial areas), and immunodeficiency related. These types have identical morphological, immunophenotypic, and genetic characters. The endemic variant is highly associated with areas where malaria infections are endemic and EBV is found in approximately all presented cases. The sporadic type occurs mostly throughout the rest of the world (mainly, North America and Europe) it is however, rarely associated with EBV infections. The immunodeficiency-related type is usually detected in immunocompromised individuals (HIV patients and organ-recipients)^(2)(5)(9)(10)(6)^. Burkitt’s lymphoma (BL) present aggressively with tumors arising from Waldeyer’s ring and/or the abdomen and more highly associated with central nervous system (CNS) and bone marrow involvement ^(3)(11)^.

*MYC* gene (MIM ID 190080) is an important proto-oncogene transcriptional factor encodes a nuclear phosphoprotein for intracellular processes. For example, regulation of cell cycle, histone acetylation and ribosomal biogenesis, programmed cell death and cellular changes. It is noticeable that the gene is increasingly expressed in several hematological malignancies including aggressive form of the B-cell lymphoma, particularly, Burkitt’s lymphoma (BL) which is an example for *MYC* gene overexpression resulting from translocation of chromosomes involving the c-*MYC* gene ^(12-18)^. *MYC* gene is the main regulator of Burkitt’s lymphoma. However, c-*MYC* gene translocation can also present less often in other types of B-cell lymphoma with other co-associated genes in lymphomagenesis such as, transcriptional factor TCF-3 and phosphoinositide-3-kinase (PI3K) pathway activator ^(16)(19)(20)^. The normal c-*MYC* gene is encoded in three exons interrupted by two large non-coding sequences. The structure and nature of the translocated gene’s linkage to the immunoglobulin locus and the presence of two c-*MYC* promoters thus resulting in two long sequences raise high chances for the activation of the oncogene.^(18)(21)^

Aberrations of the c-*MYC* proto-oncogene located in chromosome 8 are the main features of BL and other types of aggressive B-cell lymphoma.^(13)(22)^ Partial joining of chromosome 8 to either one of the chromosomes (2, 14, or 22) that bear immunoglobulin genes will deregulate *MYC* gene once translocated ^(23)(24)^. The t(8;14)(q24.1;q32) [Ig Heavy chain], and its variants – the t(2;8)(p12;q24.1) [Kappa], and t(8;22)(q24.1; q11.2)[lambda] are associated with B-cell lymphoma and result in *MYC*/immunoglobulin (*IG*) gene rearrangement. ^(9,25-30)^

This study aims to detect single nucleotide polymorphisms (SNPs) in *MYC* gene associated with formation of BL, and to confirm or exclude most reported SNPs relationship with the disease, and aspires to shed light on newly detected mutations associated with the disorder if present.

In this paper, data was obtained from national center of biotechnology information (NCBI) and big data analysis was performed through several translational tools which work using different strategies to introduce results that confirm or exclude previous studies’ findings.

## Materials and Methods

A total number of 2868 nsSNPs were obtained from the national center for biotechnology information (NCBI) on April 2018. 286 SNPs were *Homo sapiens* to which the analysis was performed.

## Mutation effect on proteins

### Based on conservation

SNPs were submitted to Sorting Tolerant from Intolerant sever **SIFT** to sort SNPs according to their effect on the protein sequence. The server divides the result into [deleterious and tolerated] according to score 0-1 respectively. SNPs with scores of 0.0 are considered to be deleterious and are further analyzed to identify the damaging ones, while SNPs with score of >0.1 are considered tolerated and not further analyzed. (Available at: http://sift.jcvi.org/) (Fig1)^(31-35)^

**Figure (1):**
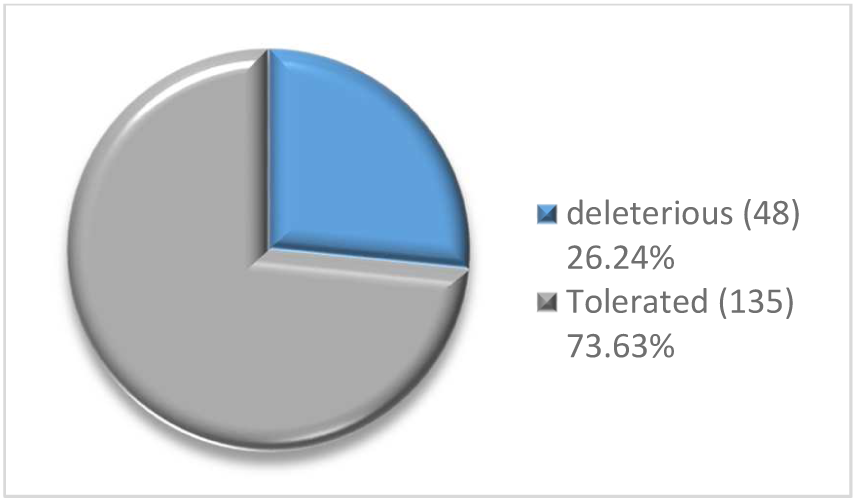
Initial analysis of single nucleotide polymorphisms SNPs using SIFT server showing numbers and percentages in two ways, Deleterious (affects protein structure and function) and Tolerated (does not alter either protein structure or functionality).

### UniProt server

Is an online website that is used to obtain a protein sequence simply by submitting its SNPs ID (rs number) or by submitting the ensemble protein ID provided by the SIFT server. The result appears as entriez and the website gives each protein a Uniprot accession number. Sequence then can be obtained in detailed manner or FASTA format which necessary for the following steps. (available at: https://www.uniprot.org) ^(36)^

### Based on structure and function

A total number of 48 deleterious single nucleotide polymorphisms SNPs were submitted to Polymorphism Phenotype v2 **Polyphen-2** server. The server differentiates SNPs according to their ability to cause a damage to the protein functionality into [probably damaging] that is the most disease causing with a score of 0.7-1, [possibly damaging] with a less disease causing ability with a score of 0.5-0.8 and [benign] which does not alter protein functions with a score of 0-0.4. (Available at: http://genetics.bwh.harvard.edu/pph2/) (Fig 2) ^(37-40)^

**Figure (2):**
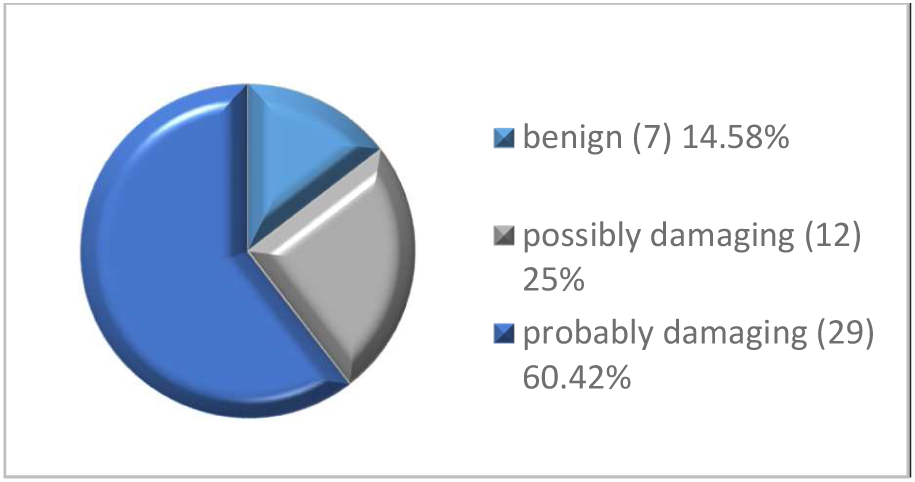
Showing numbers and percentages of single nucleotide polymorphisms SNPs and their possible results of the PolyPhen-2 server.

### Based on function

The BioFolD **SNPs&Go** server is a conformational step to ensure the disease relationship with the SNPs studied. It gives three different results based on three different analytical algorithms, panther result, PHD-SNP result, and SNPs&GO result. Each one of the results is composed of three main parts, the prediction which decides whether the mutation is neutral or disease related, reliability index (RI), and disease probability (if >0.5 mutation is disease causing). This step has not been carried out for SNPs with a result of Benign in the Polyphen-2 server. (Available at: http://snps.biofold.org/snps-and-go/snps-and-go.html) (Fig 3)^(40)(41)^

**Figure (3):**
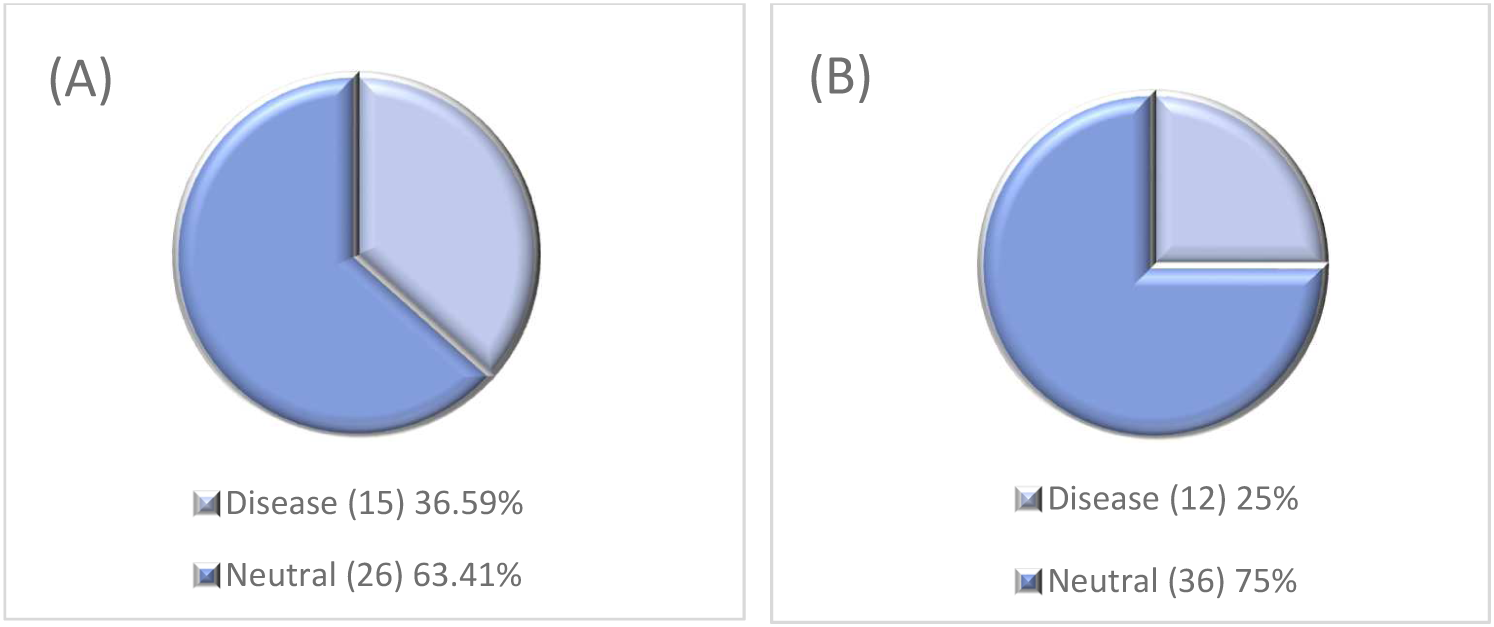
Result of single nucleotide polymorphisms SNPs analysis using SNPs&GO server **(A)** shows numbers and percentages of SNPs which are neutral (non-disease causing) or Disease causing according to SNPs&GO algorithm. **(B)** Shows numbers and percentages of SNPs which are either neutral or disease related according to PHD-SNP algorithm.

### Protein stability analysis

**I-mutant** server is another BioFolD server which is used for the analyzed SNPs to study protein stability changes caused by different mutations and to decide whether the mutation stabilizes or destabilizes the protein. The result appears in three parts, prediction part, reliability index RI, and a DDG value which is calculated from the unfolding Gibbs free energy value of the mutant type minus the unfolding Gibbs free energy value of the wild type (Kcal/mol).(Available at: http://folding.biofold.org/cgi-bin/i-mutant2.0.cgi) (Fig 4)

**Figure (4):**
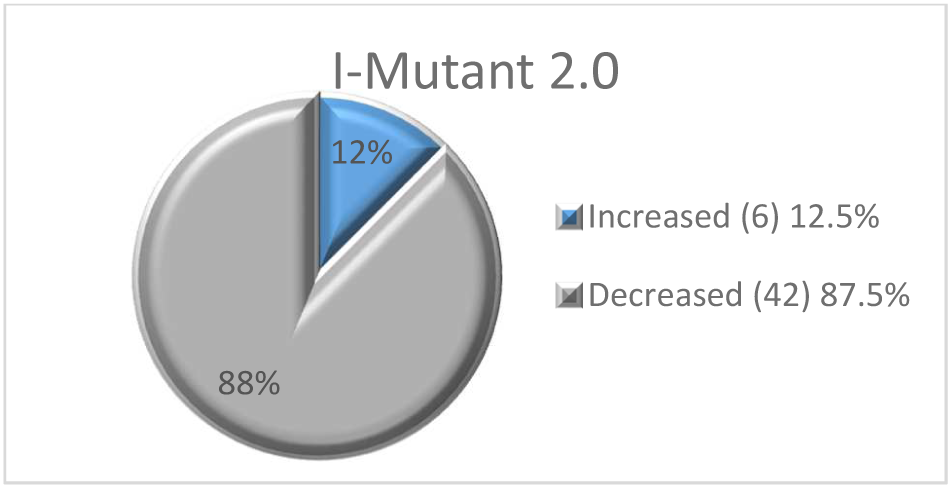
Number and percentages of single nucleotide polymorphisms SNPs analyzed using I-Mutant 2.0 server which reads the effect of a specific point mutation on protein stability

**Figure (5):**
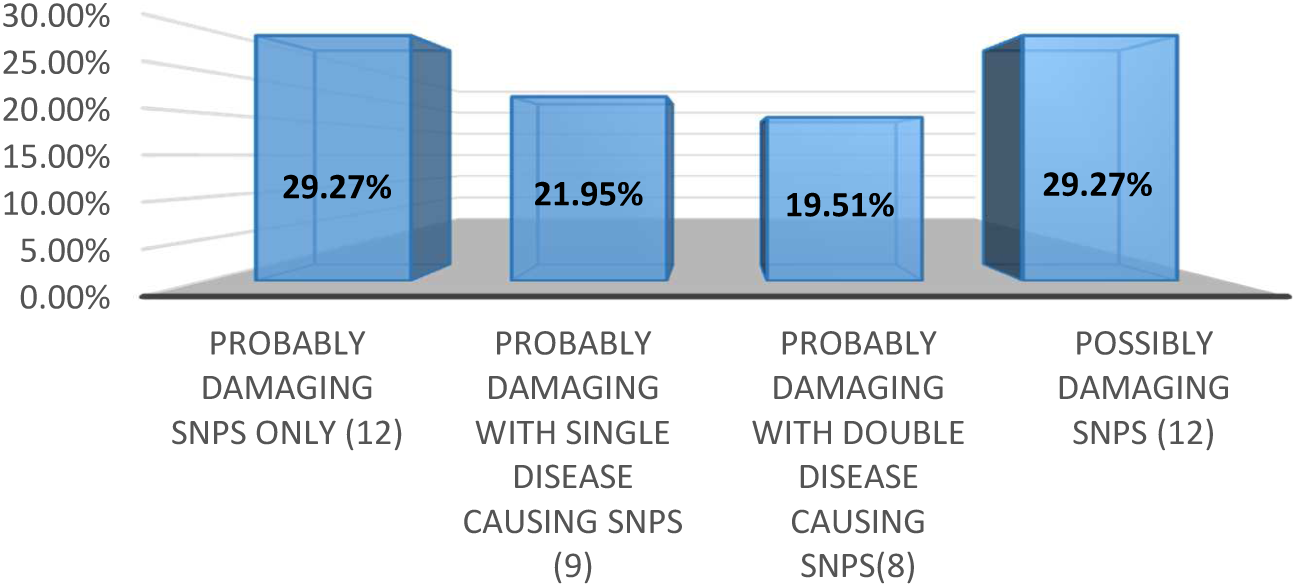
Chart showing numbers and percentages of SNPs response to polyPhen-2, SNPs&GO, PHD-SNP servers’ altogether.

**Figure (6):**
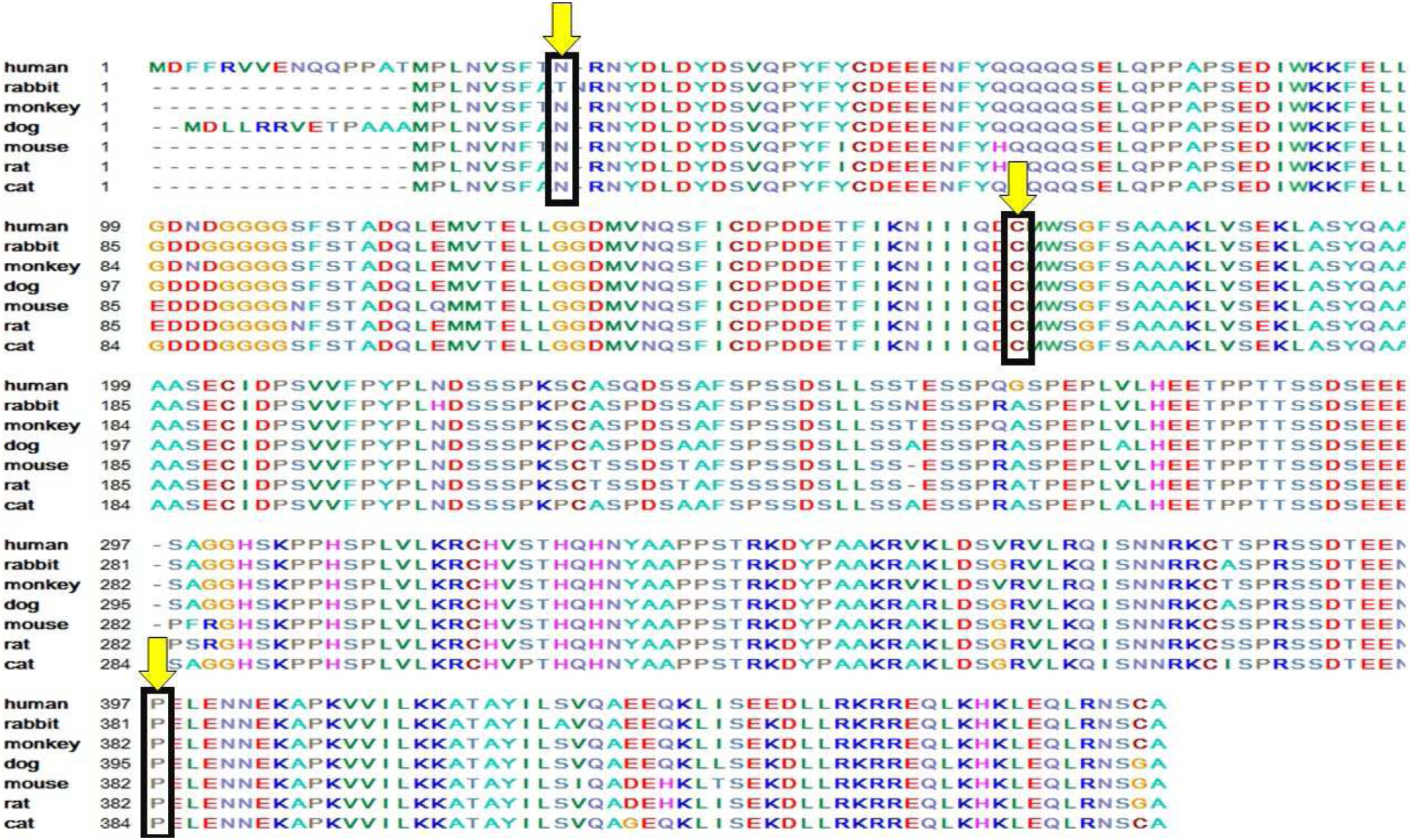
Multiple sequence alignment using Bioedit software visualizes sites of three mutations as an example are located at a highly conservative regions of C-MYC protein. **(A):** rs4645959 (protein ID ENSP00000367207- Uniprot ID P01106) mutation of Asparagine into Serine at position 26 (N26S). **(B):** rs150308400 (Protein ID ENSP00000259523 – Uniprot ID A0A0B4J1R1) mutation of Cysteine to Tyrosine at position 133 (C133Y). **(C):** rs141095253 (protein ID ENSP00000367207- Uniprot ID p01106) mutation of Proline to Leucine at position 397 (P397L).

### Structural effect of SNPs

**Mutpred2** is a tool that detects the effect of missense mutations on protein structure by providing it with the protein sequence in a FASTA format, mutation position, wild and mutant types. The result contains Mutpred2 score which is the probability that the amino acid substitution is pathogenic [>0.50 is considered pathogenic], affected PROSITE and ELM motifs, and a table with the structural changes occurred whether loss/gain of a helix, strand or loop. Also, gives other structural changes information if any and a P-value. (Table 3) (Available at: http://mutpred2.mutdb.org/index.html) ^(42-44)^

**Table (1):** SNPs result using SIFT server and PolyPhen-2 server

| SNP ID | Amino acid change | SIFT |  | polyPhen-2 |  |
| --- | --- | --- | --- | --- | --- |
|  |  | score | Prediction | value | prediction |
| rs4645959 | N25S | 0.007 | Deleterious | 1 | probably damaging |
| rs4645959 | N26S | 0.045 | Deleterious | 1 | probably damaging |
| rs4645959 | N25S | 0.045 | Deleterious | 1 | probably damaging |
| rs4645960 | G174R | 0.011 | Deleterious | 1 | probably damaging |
| rs4645960 | G160R | 0.015 | Deleterious | 1 | possibly damaging |
| rs4645960 | G175R | 0.022 | Deleterious | 1 | possibly damaging |
| rs4645960 | G174R | 0.022 | Deleterious | 1 | possibly damaging |
| rs28933407 | P71S | 0.018 | Deleterious | 1 | probably damaging |
| rs28933407 | P57S | 0.021 | Deleterious | 1 | probably damaging |
| rs28933407 | P72S | 0.043 | Deleterious | 0.394 | benign |
| rs28933407 | P71S | 0.044 | Deleterious | 1 | probably damaging |
| rs112602073 | P220S | 0.049 | Deleterious | 0.376 | benign |
| rs112602073 | P219S | 0.049 | Deleterious | 0.778 | possibly damaging |
| rs114570780 | Y32H | 0 | Deleterious | 0.794 | possibly damaging |
| rs114570780 | Y47H | 0 | Deleterious | 0.996 | probably damaging |
| rs114570780 | Y46H | 0 | Deleterious | 0.729 | possibly damaging |
| rs121918685 | P59A | 0 | Deleterious | 1 | probably damaging |
| rs121918685 | P74A | 0 | Deleterious | 1 | probably damaging |
| rs121918685 | P73A | 0 | Deleterious | 1 | probably damaging |
| rs121918685 | P73A | 0.003 | Deleterious | 0.997 | probably damaging |
| rs137906262 | L158I | 0.009 | Deleterious | 0.987 | probably damaging |
| rs137906262 | L144I | 0.019 | Deleterious | 0.987 | probably damaging |
| rs137906262 | L159I | 0.025 | Deleterious | 0.999 | probably damaging |
| rs137906262 | L158I | 0.025 | Deleterious | 0.999 | probably damaging |
| rs141095253 | P397L | 0.029 | Deleterious | 1 | probably damaging |
| rs141095253 | P396L | 0.029 | Deleterious | 0.999 | probably damaging |
| rs145561065 | L434F | 0.017 | Deleterious | 0.959 | probably damaging |
| rs145561065 | L433F | 0.018 | Deleterious | 0.326 | benign |
| rs148228388 | A76E | 0.022 | Deleterious | 0.736 | possibly damaging |
| rs148228388 | A90E | 0.032 | Deleterious | 0.853 | probably damaging |
| rs148863193 | R439L | 0.026 | Deleterious | 0.6 | possibly damaging |
| rs148863193 | R438L | 0.027 | Deleterious | 0.974 | probably damaging |
| rs150308400 | C133Y | 0 | Deleterious | 1 | probably damaging |
| rs150308400 | C148Y | 0 | Deleterious | 0.991 | probably damaging |
| rs150308400 | C147Y | 0 | Deleterious | 1 | probably damaging |
| rs150308400 | C147Y | 0 | Deleterious | 1 | probably damaging |
| rs200431478 | S362F | 0.003 | Deleterious | 0.998 | probably damaging |
| rs200431478 | S361F | 0.003 | Deleterious | 0.999 | probably damaging |
| rs370338821 | S213F | 0.035 | Deleterious | 0.848 | possibly damaging |
| **rs370767192** | H444Y | 0.014 | Deleterious | 0.168 | benign |
| **rs370767192** | H443Y | 0.014 | Deleterious | 0.241 | benign |
| **rs375812644** | L28V | 0.012 | Deleterious | 0.951 | possibly damaging |
| **rs375812644** | L29V | 0.024 | Deleterious | 0.629 | possibly damaging |
| **rs375812644** | L28V | 0.024 | Deleterious | 0.951 | possibly damaging |
| **rs375812644** | L14V | 0.026 | Deleterious | 0.964 | probably damaging |
| **rs139294902** | P11R | 0 | Deleterious | 0.037 | benign |
| **rs143501729** | A454E | 0.009 | Deleterious | 0.003 | benign |
| **rs143501729** | A453E | 0.009 | Deleterious | 1 | probably damaging |

**Table (2):** single nucleotide polymorphisms SNPs using SNPs&go and I-mutant 2.0 servers

| SNP ID | Amino acid change | SNPs&GO |  |  | PHD-SNP |  |  | I-mutant 2.0 |  |
| --- | --- | --- | --- | --- | --- | --- | --- | --- | --- |
|  |  | Value | prediction | RI | value | prediction | RI | prediction | RI |
| **rs4645959** | N25S | 0.499 | neutral | 0 | 0.662 | disease | 3 | decreased | 9 |
| **rs4645959** | N26S | 0.251 | disease | 0 | 0.691 | disease | 4 | decreased | 9 |
| **rs4645959** | N25S | 0.521 | disease | 0 | 0.691 | disease | 4 | decreased | 9 |
| **rs4645960** | G174R | 0.033 | neutral | 9 | 0.205 | neutral | 6 | decreased | 7 |
| **rs4645960** | G160R | 0.024 | neutral | 10 | 0.176 | neutral | 6 | decreased | 7 |
| **rs4645960** | G175R | 0.028 | neutral | 9 | 0.217 | neutral | 6 | decreased | 7 |
| **rs4645960** | G174R | 0.028 | neutral | 9 | 0.218 | neutral | 6 | decreased | 7 |
| **rs28933407** | P71S | 0.663 | disease | 3 | 0.356 | neutral | 3 | decreased | 8 |
| **rs28933407** | P57S | 0.666 | disease | 3 | 0.361 | neutral | 9 | decreased | 9 |
| **rs28933407** | P71S | 0.711 | disease | 4 | 0.451 | neutral | 9 | decreased | 9 |
| **rs112602073** | P219S | 0.026 | neutral | 9 | 0.112 | neutral | 7 | decreased | 9 |
| **rs114570780** | Y32H | 0.056 | neutral | 9 | 0.36 | neutral | 7 | decreased | 1 |
| **rs114570780** | Y47H | 0.062 | neutral | 9 | 0.368 | neutral | 7 | decreased | 9 |
| **rs114570780** | Y46H | 0.062 | neutral | 9 | 0.374 | neutral | 8 | decreased | 9 |
| **rs121918685** | P59A | 0.575 | disease | 9 | 0.374 | neutral | 3 | decreased | 8 |
| **rs121918685** | P74A | 0.606 | disease | 1 | 0.239 | neutral | 5 | decreased | 5 |
| **rs121918685** | P73A | 0.606 | disease | 2 | 0.289 | neutral | 4 | decreased | 5 |
| **rs121918685** | P73A | 0.577 | disease | 2 | 0.289 | neutral | 4 | decreased | 5 |
| **rs137906262** | L158I | 0.15 | neutral | 2 | 0.239 | neutral | 5 | decreased | 5 |
| **rs137906262** | L144I | 0.148 | neutral | 7 | 0.433 | neutral | 1 | decreased | 6 |
| **rs137906262** | L159I | 0.151 | neutral | 7 | 0.435 | neutral | 1 | decreased | 6 |
| **rs137906262** | L158I | 0.151 | neutral | 7 | 0.461 | neutral | 1 | decreased | 6 |
| **rs141095253** | P397L | 0.597 | disease | 2 | 0.607 | disease | 2 | increased | 0 |
| **rs141095253** | P396L | 0.597 | disease | 2 | 0.607 | disease | 2 | increased | 0 |
| **rs145561065** | L434F | 0.085 | neutral | 8 | 0.057 | neutral | 7 | decreased | 5 |
| **rs148228388** | A76E | 0.088 | neutral | 8 | 0.33 | neutral | 3 | decreased | 2 |
| **rs148228388** | A90E | 0.107 | neutral | 8 | 0.441 | neutral | 1 | decreased | 2 |
| **rs148863193** | R439L | 0.056 | neutral | 9 | 0.198 | neutral | 6 | decreased | 7 |
| **rs148863193** | R438L | 0.056 | neutral | 9 | 0.198 | neutral | 6 | decreased | 7 |
| **rs150308400** | C133Y | 0.87 | disease | 7 | 0.925 | disease | 8 | decreased | 4 |
| rs150308400 | C148Y | 0.879 | disease | 8 | 0.939 | disease | 9 | decreased | 4 |
| rs150308400 | C147Y | 0.869 | disease | 7 | 0.923 | disease | 8 | decreased | 4 |
| rs150308400 | C147Y | 0.879 | disease | 8 | 0.939 | disease | 9 | decreased | 4 |
| rs200431478 | S362F | 0.187 | neutral | 6 | 0.322 | neutral | 4 | increased | 5 |
| rs200431478 | S361F | 0.071 | neutral | 9 | 0.176 | neutral | 6 | increased | 5 |
| rs370338821 | S213F | 0.024 | neutral | 10 | 0.079 | neutral | 8 | decreased | 1 |
| rs375812644 | L28V | 0.23 | neutral | 5 | 0.499 | neutral | 0 | increased | 0 |
| rs375812644 | L29V | 0.244 | neutral | 5 | 0.519 | disease | 0 | decreased | 7 |
| rs375812644 | L28V | 0.244 | neutral | 5 | 0.518 | disease | 0 | increased | 0 |
| rs375812644 | L14V | 0.234 | neutral | 5 | 0.511 | disease | 0 | decreased | 3 |
| rs143501729 | A453E | 0.05 | neutral | 9 | 0.188 | Neutral | 6 | decreased | 4 |

**Table (3):** showing structural effects of mutations based on MutPred2 tool

| SNP ID | UniProt ID | Amino acid change | Mutpred2 score | PROSITE/ELM motifs |
| --- | --- | --- | --- | --- |
| rs4645959 | H0YBT0 | N25S | 0.326 | - |
| rs4645959 | P01106 | N26S | 0.313 | - |
| rs141095253 | H0YBT0 | P396L | 0.597 | ELME000333 |
| rs141095253 | P01106 | P397L | 0.611 | ELME000333 |
| rs150308400 | A0A0B4J1R1 | C133Y | 0.784 | None |
| rs150308400 | H0YBG3 | C147Y | 0.866 | None |
| rs150308400 | H0YBG0 | C147Y | - | - |
| rs150308400 | P01106 | C148Y | 0.823 | None |

### Homology modeling

**Raptor X** server is a software for protein structure and function prediction developed by XU group by a given protein sequence the server can predict protein secondary and tertiary structures along with other functional information by running multiple sequence analysis. Result comes in cartooned 3D image of the predicted protein along with a Protein Data Bank PDB ID of the protein with the highest identities. (Available at: http://raptorx.uchicago.edu/)^(45-47)^

### Visualization of protein

Using **Chimera 1.8** software which is a product of University of California, San Francisco UCSF for visualization and editing of the three dimensional structure of the c-MYC protein (available at: http://www.cgl.ucsf.edu/chimera) (fig 7)^(48)^

**Figure (7):**
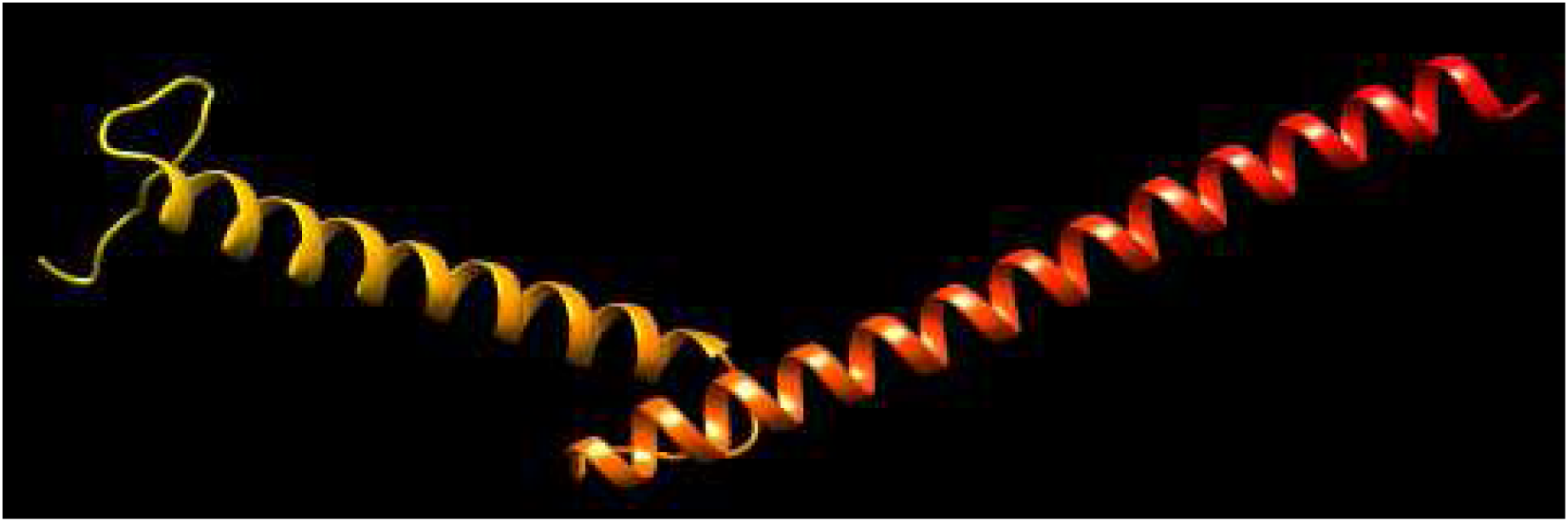
Partial 3D structure of the c-MYC protein showing alpha and beta chains of the protein using UCSF **Chimera** software (version 1.8)

### Taxonomy sequences

**NCBI blast** server was used to obtain *Homo sapiens* (human) family sequences to align the sequences of these orders. Six sequences of *Mus musculus* (mouse) [Accession: AAB59728.1], *Rattus norvegicus* (rat) [Accession: AAQ57167.1], *Oryctolagus cuniculus* (rabbit) [Accession: NP_001306504.1], *Canis lupus* (dog) [Accession: NP_001003246.2], *Macaca mulatta* (monkey) [Accession: NP_001136345.1], *Felis catus* (cat) [Accession: NP_001166917.1]. (Available at: https://blast.ncbi.nlm.nih.gov/Blast.cgi) ^(49-51)^

## Multiple sequence alignment

Using **Bioedit** software produced by **Clustal W** to view areas of high conservation between different proteins of the same family. Mutations located in highly conservative regions are most likely to cause a change in protein structure and function. The software is used on SNPs which scored Deleterious, probably damaging and double disease causing by SIFT, PolyPhen-2 and SNPs&go servers, respectively. Which are eight single nucleotide polymorphisms SNPs.^(52)^

## Results

## Discussion and Conclusion

Eight SNPs (rs4645959, rs4645959, rs141095253, rs141095253, rs150308400, rs150308400, rs150308400, rs150308400) were found to be the most disease causing SNPs (which has higher probability of altering protein structure and function thus; causing disease). Probably damaging with double disease causing result SNPs are more likely to be involved in formation of disease, while Possibly damaging SNPs are the least likely to cause disease. Moderate probability of disease association for SNPs with probably damaging only, or probably damaging with single disease causing SNPs.

SNP rs4645959 showed mutation of Aspargine into Serine at position 25 in protein isoform-1 (Uniprot accession H0YBT0) and additional SNP rs141095253 conversion of Proline into Leucine at position 396 by protein Isoform-1 (Uniprot accession H0YBT0) were found to be probably damaging, double-disease causing by PolyPhen-2, SNPs&GO and PHD-SNP, respectively. These mutations have not been reported before in previous studies regarding c-*MYC* gene analysis. In addition, they were also located in a highly conserved regions of the protein sequence.

A study by Mamoona Noreen in 2015 stated that SNP rs4645959 mutation of Aspargine into Serine at position 26 in protein Isoform-2 (protein accession P01106) was found to be benign in contrast to result obtained in this study that the same SNP was deleterious, and disease causing by PolyPhen-2, SNPs&GO respectively. at further analysis by multiple sequence alignment the mutation was located in a highly conservative region indicating the major effect of the mutation on protein’s structure and functionality^(28)^.

Mamoona Noreen’s study of 2015 also found rs141095253 mutation of proline into Leucine at position 397 by protein Isoform-2 (Uniprot accession P01106) to be probably damaging by PolyPhen-2 server, this study confirms the previous result and increases certainty by having double-disease causing prediction by SNPs&GO and PHD-SNP and through location of mutation in highly conserved region. However, differs from cited paper for having a deleterious result in SIFT server ^(28)^.

in 2016 a study by Afra Abd Elhamid stated that SNPs rs150308400 with conversion of Cysteine into Tyrosine at positions 133, 147 (two protein sequences), and 148 by protein Isoform-1 (Uniprot accessions A0A0B4J1R1, H0YBG3, H0YBG0, P01106 respectively) had given high disease prediction probability based on their results on SIFT, PolyPhen-2, and PHD-SNP servers. Which matches results obtained regarding these SNPs in this study ^(28)(40)^.

Through the use of Mutpred2 on the eight disease causing SNPs, two out of them were considered as non-pathogenic SNPs. rs4645959 (N25S) with Mutpred2 score of 0.326 and rs4645959 (N26S) with Mutpred2 score of 0.313. rs141095253 (P396L) and rs141095253 (P397L) predicted to have the same effect on protein which gained a helix and relative solvent accessibility in the secondary protein structure with a (P-value of 0.28-0.23 and 0.29-0.25 respectively). Both SNPs also affected ELM000333. Mutation with rs150308400 (C133Y) was predicted to alter protein structure by adding a strand to it and increasing sulfation of Cysteine at position 133 (P-value 0.27-0.02). rs150308400 (C147Y) with protein accession of HOYBG3 mutation affected the protein in two ways, the first was the loss of the disulfide bond in the cysteine at position 147, while the other was through extra sulfation of cysteine of the same position (P-value 0.4-0.02). SNP rs150308400 (C148Y) affected the MYC protein by gain of a strand, loop and sulfation of cysteine at position 148 (P-value 0.29-0.27-0.02, respectively).

SNP rs4645959 (N25S) and SNP rs141095253 (P396L) were found to be disease causing according to SIFT, PolyPhen-2, SNPs&GO and PHD-SNP servers and have not been reported in other studies previously. We recommend further in vivo and in vitro analysis for the previously mentioned SNPs to assess their disease particular relationship.

Limitations regarding this study, included being a fully computational research. Such studies, should be conducted in vivo and in vitro besides being conducted with bioinsilico technology. Another limitation was the inability to obtain complete protein structure due to failure of retrieving the full accessible sequence from homology modeler Raptor X and Chimera softwares. Due to continuous update of databases MYC protein with UniProt accession of HOYBG0 was removed from the database and failed to obtain the complete sequence for Mutpred2 prediction.

Translation bioinformatics is a science that uses several computational tools to help study biological information which occurs in public databases (e.g. SNPs). It helps in unravelling huge numbers of genes found in gene bank and provides full informations about a desired gene sequence^(53)^. It has been very helpful in studies of gene’s expression and abnormality that may or may not have a relationship to certain disease causing. So, they facilitate detection, monitoring and observing disease related abnormalities. Some of the applications of translation bioinformatics which are becoming the headlines of genetic diseases, are using them as diagnostic tools. Such usage is made through, SNPS genotyping arrays which uses disease causing SNPs reported in database as diagnostic markers for specific disorders. In conclusion, it is highly recommended to perform in vitro and in vivo analysis of the newly detected mutations (rs4645959 (N25S) isoform-1 [Uniprot H0YBT0]) and (rs141095253 P396L Isoform-1 [Uniprot H0YBT0]) and to obtain full three-dimensional structure of the MYC protein to validate the previous results.

